# Sex-dependent changes in insular cortex connectivity in a rat model of comorbid pain

**DOI:** 10.64898/2026.02.07.704080

**Authors:** Laura Ventura, Michael L. Keaser, Luis G. Hernández-Rojas, Mahadi Shahed, Hayelom K. Mekonen, Ohannes Melemedjian, Alison J. Scott, Robert K. Ernst, David A. Seminowicz, Richard J. Traub, Joyce T. Da Silva

## Abstract

Temporomandibular disorder (TMD) and irritable bowel syndrome (IBS) are two highly comorbid, nociplastic pain conditions that belong to a broader group of commonly co-occurring chronic pain conditions. Most of these chronic overlapping pain conditions (COPCs), including TMD and IBS, disproportionately affect females and are highly stress sensitive. Our previous study illustrated sex differences in brain activity during colorectal distension specific to our model of comorbid pain hypersensitivity (CPH), in which masseter muscle inflammation followed by restraint stress elicits IBS-like visceral hypersensitivity. Since insular cortex (Ins) activity increased in female CPH rats only and abnormal Ins activity has been identified in TMD and IBS patients, we sought to characterize patterns of Ins-based functional connectivity (FC) by performing functional MRI (fMRI) scans at baseline, 1 week, and 7 weeks post-injury/stress in groups of male and female Sprague-Dawley rats randomized to the following conditions: CPH, stress-induced hypersensitivity (SIH), Complete Freund’s Adjuvant (CFA)-induced masseter muscle inflammation, and naive. CPH females displayed extensive Ins FC with brain regions in the cortex and limbic system, including the thalamus. Compared to CPH males, CPH females showed robust insular-thalamo connectivity at week seven, a time point where visceral hypersensitivity and referred pain-like behavior persists in CPH females but not males. This trend is also apparent in CPH females’ week seven versus week one Ins FC, whereas CPH males tend to decrease Ins FC broadly. These findings potentially suggest sensitization within the insular-thalamo and -cortical networks in CPH females, warranting future investigation of Ins circuit involvement in comorbid pain.

## Introduction

Nociplastic pain, or pain caused by altered nociception without clear evidence of tissue damage [1], frequently presents as comorbid [64], especially with other chronic pain conditions [22,69]. The term chronic overlapping pain conditions (COPCs) encompasses those chronic pain conditions—many of which exhibit nociplastic features—that commonly co-occur. The co-aggregation of COPCs compounds individual disease burden and complicates treatment, highlighting the need for thorough characterization of the mechanisms driving the development and progression of COPCs [58]. TMD and IBS are two strongly associated COPCs, with one clinical study reporting a 6-fold greater likelihood of having IBS under TMD status compared to healthy status [61]. These pain conditions are highly susceptible to psychological stress, anxiety, and depression, which modulate pain severity [47,59,62,72,75]. Many COPCs, including TMD and IBS, exhibit greater prevalence and severity in females [3,12,27,56,66], a population already overrepresented in comorbid pain and mental health disorders [2,77]. We previously developed a rodent model of comorbid pain hypersensitivity (CPH) in which stress following induction of TMD-like pain produces visceral hypersensitivity resembling that of IBS [80]. We found that persistent visceral hypersensitivity is dependent on stress during pre-existing orofacial pain [80], with female CPH rats exhibiting more profound visceral hypersensitivity compared to CPH males [19]. Moreover, female rats without orofacial pain (stress-only) exhibit transient visceral hypersensitivity, and their male counterparts exhibit even shorter-duration visceral hypersensitivity [14,44,45,80]. Further, the naive and orofacial pain conditions in female rats produce no visceral hypersensitivity [14,44,80]. Psychological stress can factor into the development and exacerbation of TMD and IBS and may contribute to their overlap and female predominance; however, the brain mechanisms underlying the development of COPCs remain largely unexplored.

Among the neural substrates linked to painful TMD, the insular cortex (Ins)—a sexually dimorphic brain region involved in pain and emotional processing [41,67,82]—has been shown to exhibit altered connectivity at rest and during noxious facial stimulation in patients with painful TMD [40]. As the Ins also participates in visceroception, Ins abnormalities have been observed in patients with visceral pain conditions such as ulcerative colitis and IBS [4,25,65], the latter exhibiting changes in Ins morphology [6,31,71], biochemical profile [9], and connectivity [33,37,73], which have been linked to pain intensity [6,16,73] and disease duration [16,71]. Psychological risk factors for TMD and IBS, such as stress, anxiety, and depression, also correspond with aberrant Ins activity [49,52,86].

Previously, we found that increased Ins activity during colorectal distention coincided with greater and longer-lasting visceral hypersensitivity in female CPH rats [19]. In the current study, we employed a seed-based correlational analysis to characterize Ins-based connectivity patterns in rats under CPH and under individual components of the CPH model that evoke shorter-duration/no hypersensitivity: stress, orofacial pain, and naive conditions [44,45]. We report increased insular-thalamo connectivity in CPH females at a time point where we previously identified persistent visceral hypersensitivity and referred pain-like behavior in CPH females only, possibly suggesting that sensitization within this network may be a female-specific mechanism promoting pain chronicity in this model. In male CPH rats, decreasing Ins connectivity with thalamic, cortical, and limbic regions over time may, then, potentially reflect male-specific silencing of pronociceptive circuits, possibly explaining pain resolution in this group.

## Materials and methods

### Animals and ethical approval

Scans were performed on male and female Sprague-Dawley rats (Envigo, 8 weeks old at time of arrival). Rats were acclimated to the animal housing facility at 22 to 23°C on a 12-hour light/dark cycle at least 7 days before experiment start. All protocols were approved by the University of Maryland Institutional Animal Care and Use Committee and conform to the Guide for use of Laboratory Animals by the International Association for the Study of Pain. CPH rats were injected bilaterally in the masseter muscle with Complete Freund’s Adjuvant (CFA, Sigma Aldrich, St. Louis, MO, F5881; 50 µl, 1:1 in saline) followed by 4 days of restraint stress, as previously described [19]. Stress-Induced Hypersensitivity (SIH) rats only underwent restraint stress. CFA rats were injected bilaterally in the masseter muscle with CFA only, and naive rats did not receive injections nor undergo restraint stress. MRI scans were performed on all rats for 3 sessions: Baseline (pre pain/stress) and 1- and 7-weeks post injury/stress. Procedure and imaging technicians were blinded to the treatment conditions.

### Comorbid Pain Hypersensitivity (CPH model)

Masseter muscles were injected with CFA [44] one day prior to initiating the restraint stress protocol: 4 days of restraint stress for 2 hours in movement-restricting tubes alternating 45° head up, head down, or flat at 15-minute intervals [19]. The day following the last stress session was designated as day one. Baseline MRI scans were collected prior to the CFA injection/restraint stress.

### fMRI acquisition parameters and analysis

Data were acquired using a Bruker BioSpec 94/30USR 9.4-T scanner (Bruker Biospin MRI GmbH, Ettlingen, Germany) and a 2×2 array coil. During scanning, rats were anesthetized at a constant mixture of 1.5% isoflurane in oxygen-enriched air. Per our previous work and that of other groups, low-dose isoflurane anesthesia has been shown to detect and generally preserve connectivity metrics, is highly reproducible between species, and does not require invasive procedures, such as intraperitoneal and subcutaneous injections in the case of medetomidine [13,30,83]. Anesthesia by definition can alter brain activity, influencing fMRI findings; however, unwanted effects are not exclusive to isoflurane. Rather, as illustrated by Paasonen et al, functional connectivity was shown to be uniquely modulated across anesthesia protocols [68]. Further, while we acknowledge the growing implementation of low-dose isoflurane in combination with medetomidine in rodent fMRI studies, the potential long-term effects of medetomidine on nociception are uncertain, as medetomidine, an α2 adrenergic receptor agonist, has demonstrated analgesic effects in rodents [70] and requires administration of atipamezole to decrease its side effects, a secondary procedure which itself may present unwanted analgesic, sedative, or negative cardiovascular effects. Finally, considering that isoflurane-only protocols are currently being employed in rigorous studies [5,78] and that restraint training for awake fMRI protocols may alter stress responses and nociceptive behavior in rodents [57], we have identified low-dose isoflurane as an appropriate anesthetic for a longitudinal fMRI study in rats undergoing a chronic pain model.

During scanning, respiration and heart rate were monitored with a small animal monitoring and gating system and software (SA Instruments, Inc, Stony Brook, NY). T2-weighted images were obtained using a 2D RARE sequence (400 × 400 matrix, 22 coronal 1-mm slice thickness, resolution 0.1 x 0.1 x 1 mm, TR 3096 ms, TE 42.2 ms). Functional MRI scans were acquired during a resting state scan for 15.5 minutes; using an echo planar imaging (EPI) sequence (TR 1511 ms, TE 24 ms, 128 × 128 matrix, resolution 0.3125 × 0.3125 × 1.0 mm, 22 coronal slices, 620 volumes per scan).

Functional datasets were preprocessed using RABIES version 0.5.1 [23] and AFNI [17] software packages. Preprocessing steps included slice-time correction, motion correction, co-registration between the functional image and its corresponding anatomical image, spatial normalization to the SIGMA rat brain template (0.4mm isotropic voxel) [7], bandpass filtering (0.009-0.2Hz), and spatial smoothing (FWHM 0.8mm). Using SPM [24], a seed-based analysis was conducted using bilateral granular insular cortex as a seed. BOLD time series contained within each voxel of the insula seed were averaged to produce an average seed time series. This time series was used as a regressor of interest at each voxel across the whole brain to reveal insula functional connectivity (FC) for each animal. Six motion parameters were also included as regressors of no interest. First-level β-contrast images representing granular insula FC were used as dependent variables in a group-level analysis measuring the effects of time (baseline, week1, and week7) and group (CPH, CFA, SIH, and NAIVE). This group-level analysis involved using the Sandwich Estimator (SwE) toolbox [32], examining group longitudinal (time), sex, and cross-sectional (group) differences. The SwE toolbox uses an ordinary least squares marginal model to generate estimates of the parameters of interest and estimate variances/covariances accounting for repeated-measures correlation associated with longitudinal data. Group-level contrast maps depicting group, time, and sex differences were cluster size corrected at p = 0.05 and combined with an individual voxel threshold of 0.05, 0.005, or 0.001. AFNI’s 3dClustSim was used to determine the minimum cluster size criteria per individual threshold. Brain regions were identified using the Waxholm atlas [51]. The Paxinos and Watson atlas (5th edition, 2004) was used as a reference for brain region abbreviations (see Table 1).

**Table 1.**
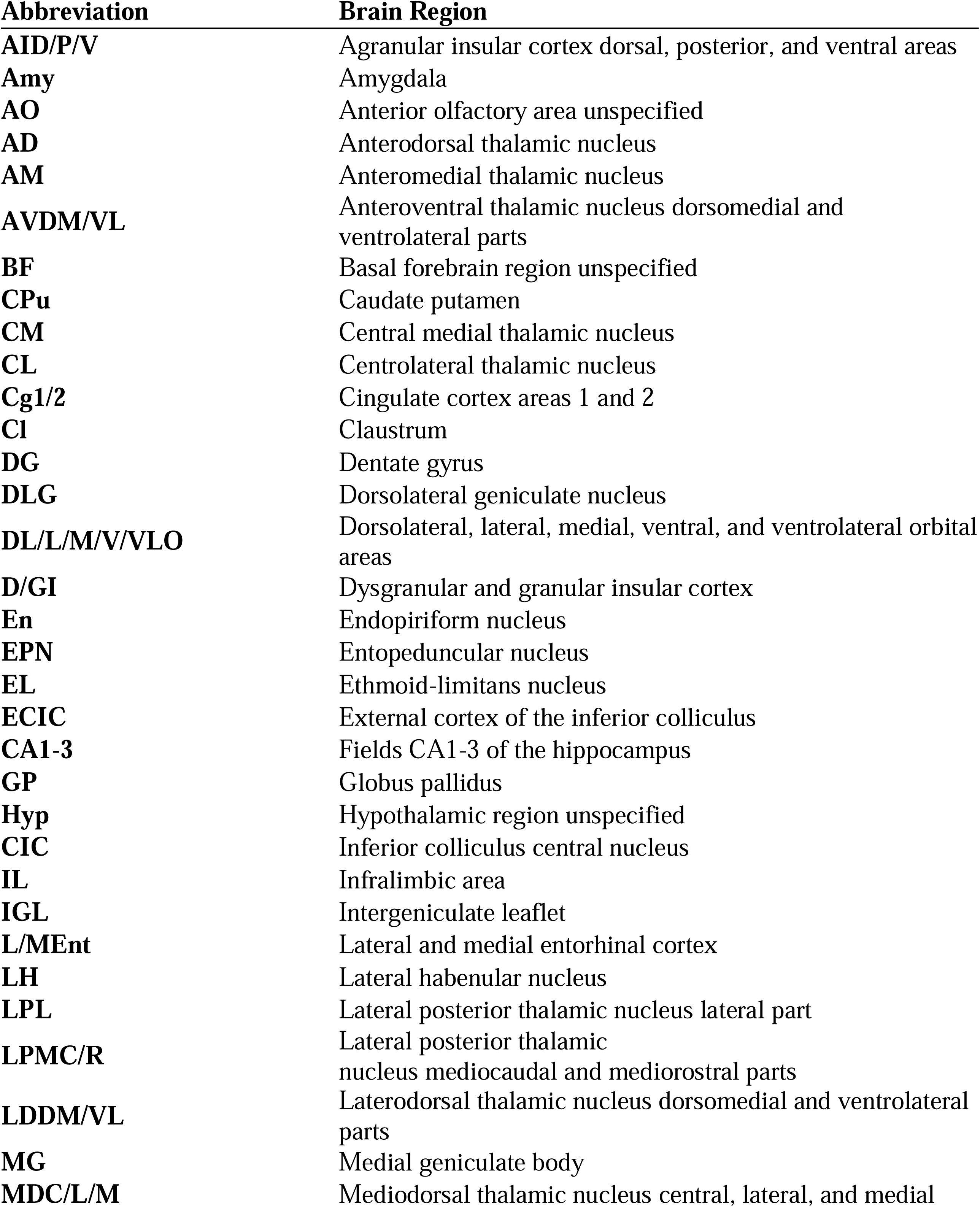
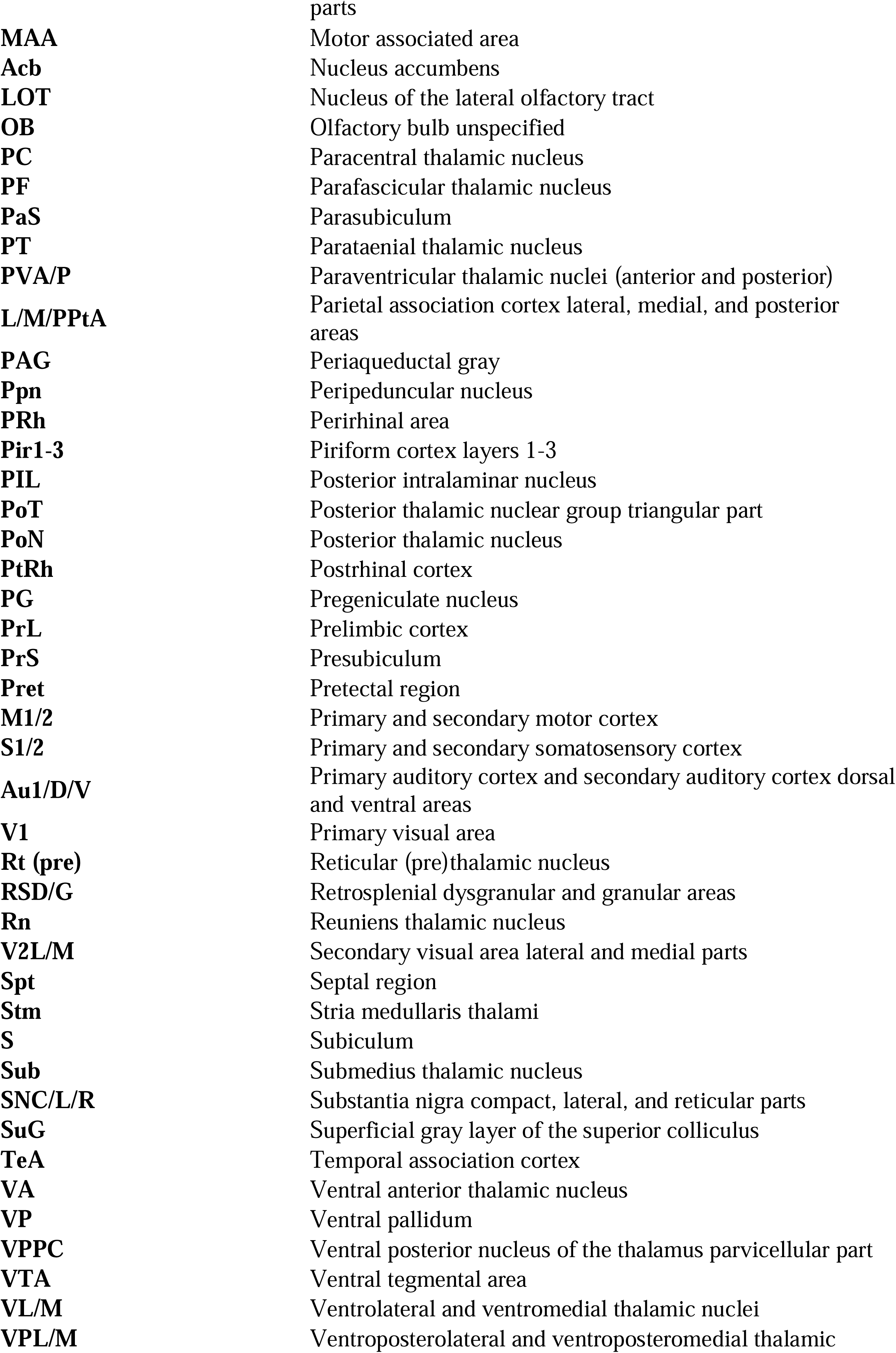
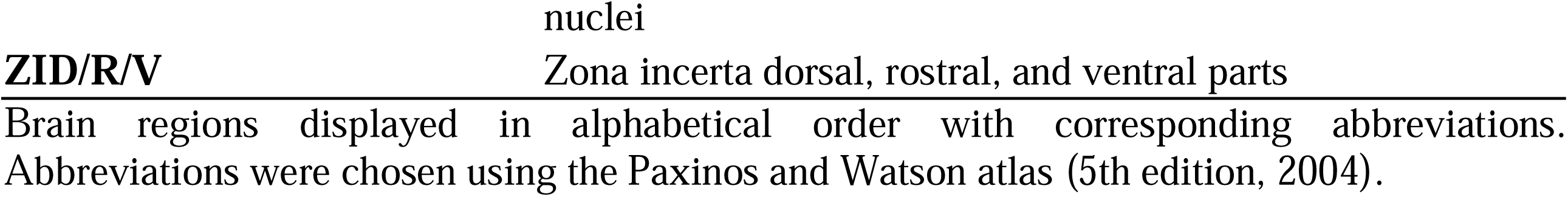
Brain regions and corresponding abbreviations.

## Results

### Insula connectivity patterns under comorbid pain, stress alone, orofacial pain, and naive conditions

We previously reported that stress induces transient visceral and orofacial hypersensitivity which is significantly prolonged if orofacial inflammation precedes the stress [44,80]. Therefore, based on our previous findings regarding increased Ins activity in the CPH model, we decided to isolate the Ins as a region of interest for a seed-based correlational analysis to elucidate potential Ins involvement in the CPH model as well as under SIH, CFA, and naive conditions (Fig. 1). Figures 2-5 show one-sample t-test maps of Ins-based resting state connectivity for male and female CPH, SIH, CFA, and naive rats, respectively, at baseline, week one, and week seven (see Supplementary Fig. 1 for all significant brain regions). Female CPH rats exhibit robust Ins FC (Fig. 2A), notably with cortical areas and thalamic nuclei, while FC with these areas in CPH males was observed to a lesser extent (Fig. 2B). Female and male SIH groups show similar FC patterns to their CPH counterparts (Fig. 3); however, Ins FC at week seven in SIH females overall exhibits less apparent FC to cortical areas and thalamic nuclei. In CFA and naive groups, less extensive Ins FC can be observed at each time point, especially progressing more caudally (Fig. 4-5). In contrast to CFA and naive groups, CPH and SIH groups, especially CPH females, show robust Ins FC at multiple brain levels, especially cortical and thalamic regions. Thus, we decided to focus on Ins FC differences between the CPH and SIH groups to investigate how prior injury leads to greater pain sequela in stressed females. Overall, CPH females tend to increase Ins FC with cortical and thalamic brain areas, suggesting that sensitization within these networks may contribute to prolonged pain-like responses in the CPH model.

**Figure 1.**
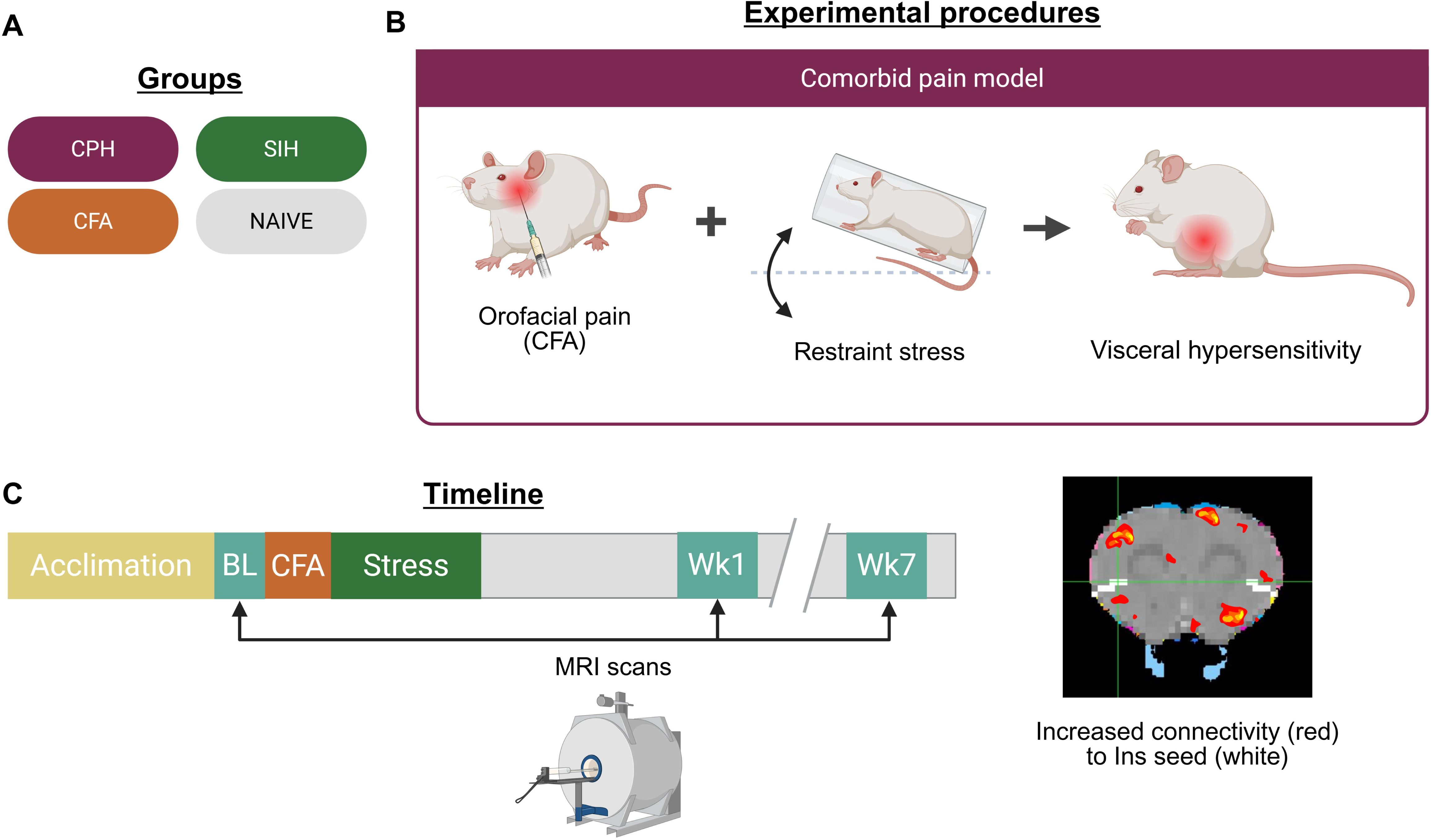
Study design. (**A**) The study consisted of the following groups: comorbid pain hypersensitivity (CPH), stress-induced hypersensitivity (SIH), Complete Freund’s Adjuvant (CFA)-induced masseter muscle inflammation (orofacial pain), and naive female (F) and male (M) rats. (**B**) The comorbid pain model consists of CFA-induced orofacial pain followed by restraint stress, which results in visceral hypersensitivity. CFA groups only undergo bilateral masseter muscle CFA injection, while SIH groups only undergo restraint stress. Naive groups receive neither CFA injection nor undergo restraint stress. (**C**) After at least 7 days of animal facility acclimation, all rats undergo a baseline (BL) MRI scan. Rats in each group then undergo the corresponding protocol (comorbid pain, stress only, CFA only, or nothing). 1 week and 7 weeks after injury/stress, all rats undergo week one (Wk1) and week seven (Wk7) scans, respectively. A seed-based correlational analysis is employed to assess Ins-based functional connectivity in all groups at each time point. Red marks on the brain image are superimposed graphics used to illustrate increased connectivity to insula (Ins) seed in white. Created in BioRender. Da Silva, J. (2025) https://BioRender.com/rna33op.

**Figure 2.**
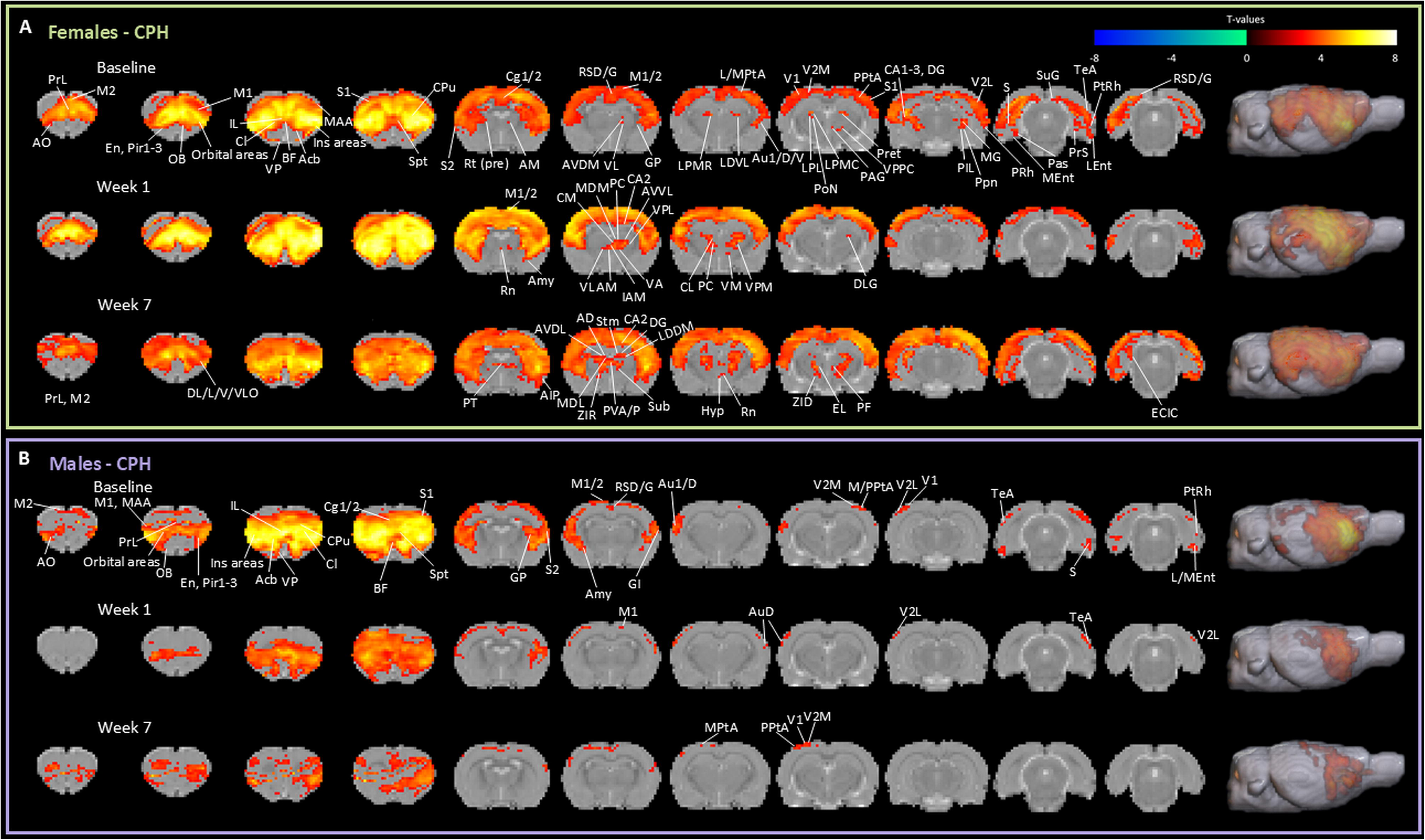
Insula-based resting state functional connectivity in CPH females and males. Illustrated brain regions show increased connectivity with Ins at baseline, week one, and week seven in CPH females (baseline: *n* = 16; week one: *n* = 18; week seven: *n* = 13) (**A**) and CPH males (baseline: *n* = 16; week one: *n* = 11; week seven: *n* = 12) (**B**). Group-level, one-sample t-test maps show significant voxels at *p* < 0.001 with cluster size correction of *p* = 0.05. See Table 1 for brain regions and corresponding abbreviations.

**Figure 3.**
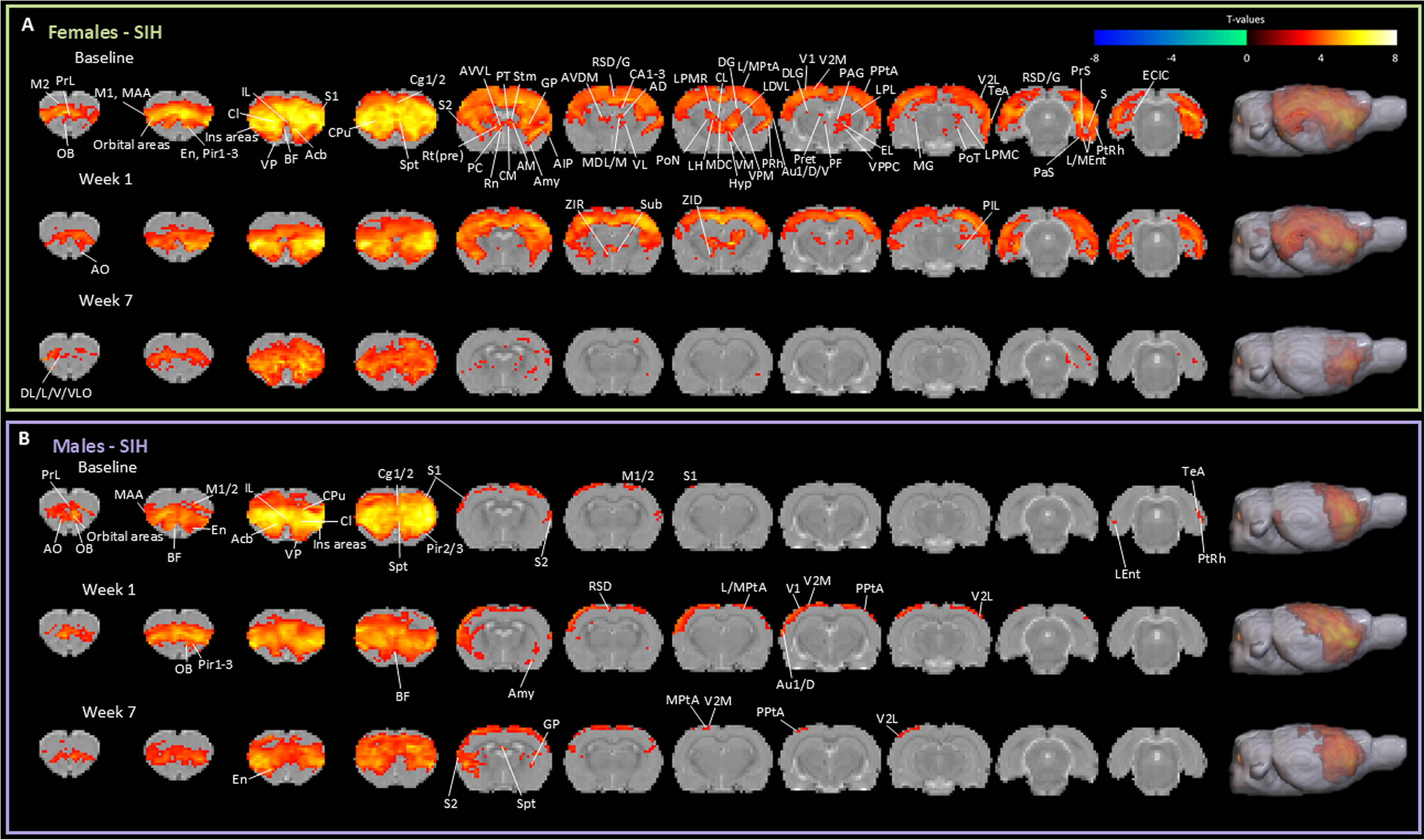
Insula-based resting state functional connectivity in SIH females and males. Illustrated brain regions show increased connectivity with Ins at baseline, week one, and week seven in SIH females (baseline: *n* = 13; week one: *n* = 15; week seven: *n* = 11) (**A**) and SIH males (baseline: *n* = 12; week one: *n* = 11; week seven: *n* = 11) (**B**). Group-level, one-sample t-test maps show significant voxels at *p* < 0.001 with cluster size correction of *p* = 0.05. See Table 1 for brain regions and corresponding abbreviations.

**Figure 4.**
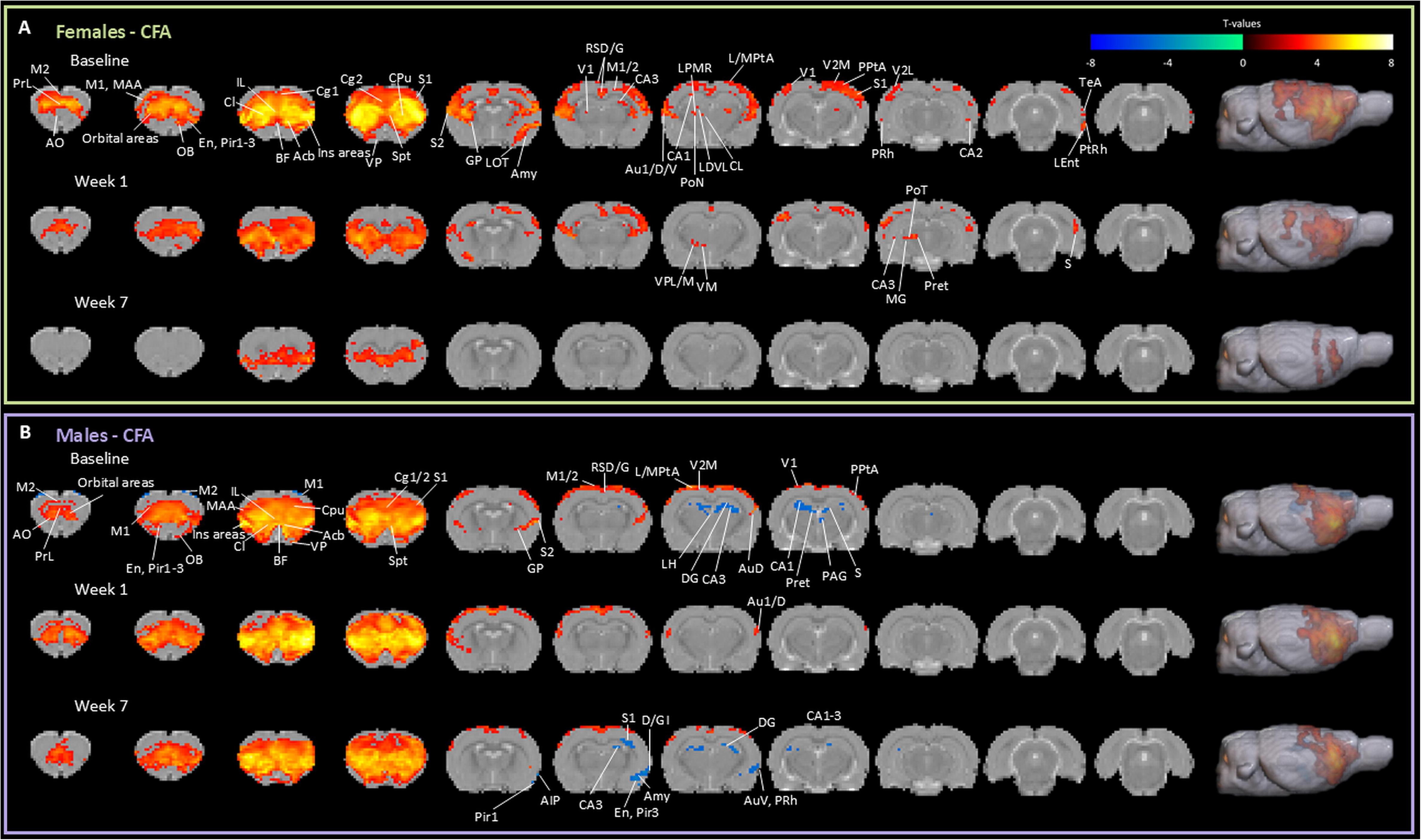
Insula-based resting state functional connectivity in CFA females and males. Illustrated brain regions show increased and decreased connectivity in warm and cold colors, respectively, with Ins at baseline, week one, and week seven in CFA females (baseline: *n* = 17; week one: *n* = 11; week seven: *n* = 8) (**A**) and CFA males (baseline: *n* = 14; week one: *n* = 13; week seven: *n* = 9) (**B**). Group-level, one-sample t-test maps show significant voxels at *p* < 0.001 with cluster size correction of *p* = 0.05. See Table 1 for brain regions and corresponding abbreviations.

**Figure 5.**
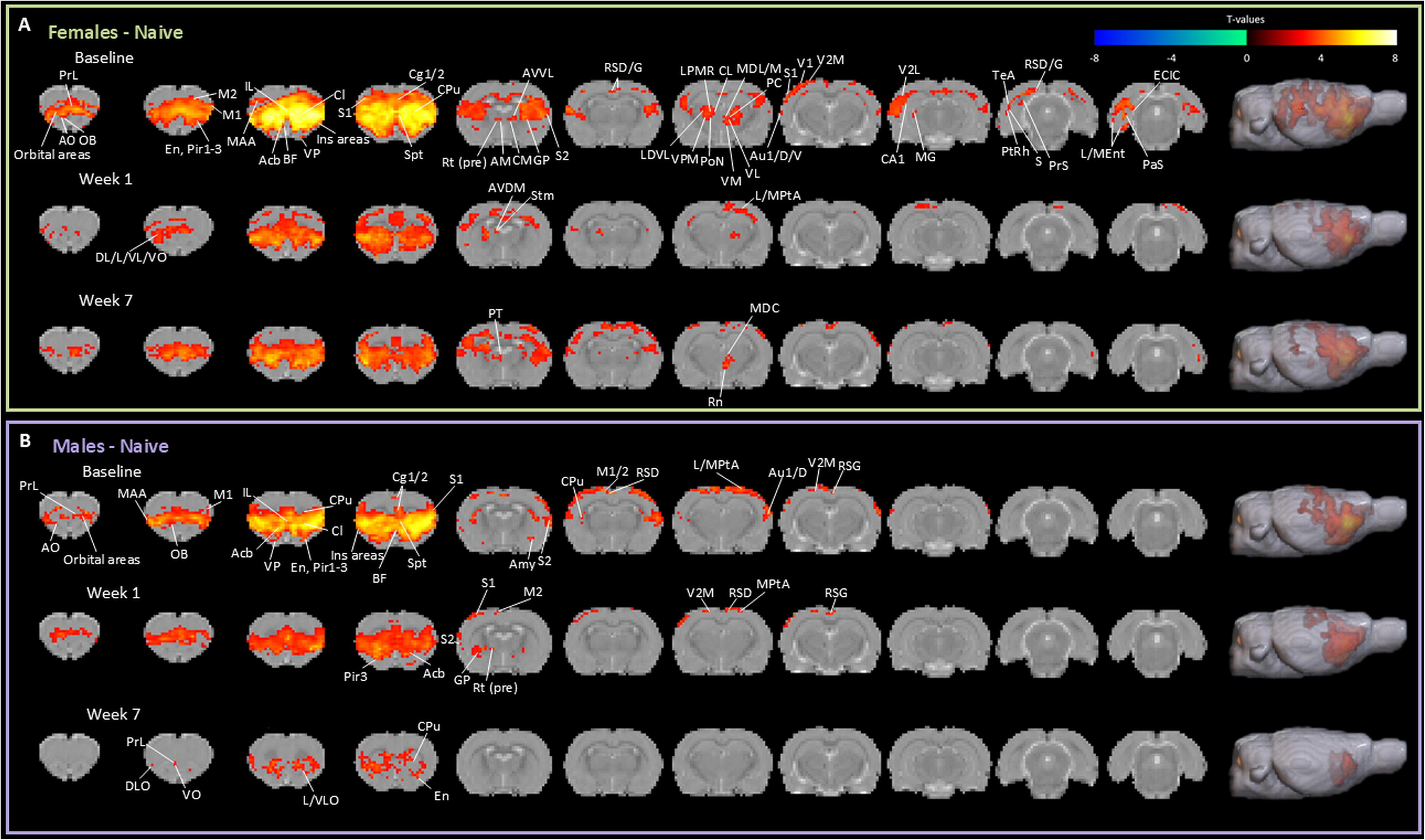
Insula-based resting state functional connectivity in naive females and males. Illustrated brain regions show increased connectivity with Ins at baseline, week one, and week seven in naive females (baseline: *n* = 17; week one: *n* = 9; week seven: *n* = 10) (**A**) and naive males (baseline: *n* = 16; week one: *n* = 7; week seven: *n* = 4) (**B**). Group-level, one-sample t-test maps show significant voxels at *p* < 0.001 with cluster size correction of *p* = 0.05. See Table 1 for brain regions and corresponding abbreviations.

### Insula connectivity differences in comorbid pain and stress alone

Our previous data shows that stress alone produces visceral hypersensitivity lasting at least 2 weeks in female rats and just days in males [44,45]. Interestingly, in female rats that undergo both CFA injection and restraint stress, visceral hypersensitivity is prolonged compared to under stress alone (SIH) [80]. Thus, we compared Ins-based FC patterns in CPH and SIH groups to better understand the brain mechanisms contributing to the maintenance of visceral hypersensitivity under prior injury and stress (see Supplementary Fig. 1 for all significant brain regions). At week one, SIH females show greater Ins FC with orbital areas (L/V/VLO) compared to CPH females, while CPH females exhibit increased Ins FC with motor areas (M1/2 and MAA), Cg1, insular areas (D/GI and AID/V), Pir3, S1/2, CPu, L/M/PPtA, and V1/2 (Fig. 6A). At week seven, during which visceral hypersensitivity remains in CPH females but not SIH females, CPH females maintain increased Ins FC with M1/2, Cg1, GI, S1/2, CPu, PtA, and V1/2 with the addition of increased Ins FC to PrL, Cg2, Cl, GP, RSD/G, Au1/D/V, S, PRh, PtRh, Ent, and TeA compared to SIH females. In contrast to the vast differences between female groups, male groups only differed at baseline (Fig. 6B), which may reflect individual variability (further discussed in limitations), as SIH and CPH females also differed modestly at baseline. These results indicate that prior injury may potentially sensitize the Ins to subsequent stress-induced neuronal alterations, engendering vast connectivity changes seen by week seven that may subserve continued visceral hypersensitivity at this time point in female CPH rats [44,80]. In contrast, male CPH and SIH groups did not differ at any time points post-stress, so we proceeded to interrogate sex differences in Ins-based connectivity within the CPH groups.

**Figure 6.**
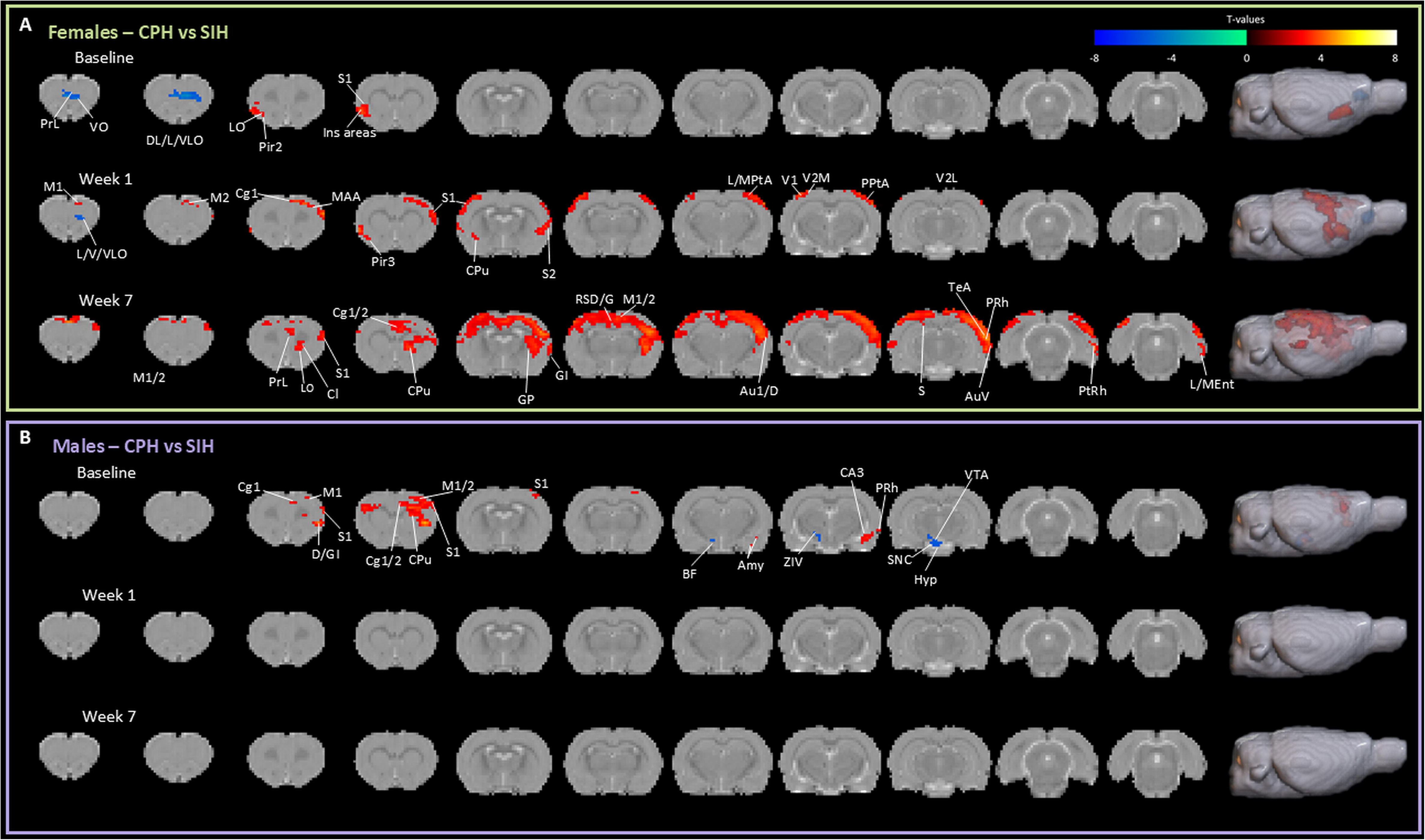
Effect of prior injury on insula-based connectivity in stressed female and male rats: Changes under comorbid pain and stress alone. Illustrated brain regions show increased and decreased connectivity in warm and cool colors, respectively, with Ins at baseline, week one, and week seven in CPH females (baseline: *n* = 16; week one: *n* = 18; week seven: *n* = 13) relative to SIH females (baseline: *n* = 13; week one: *n* = 15; week seven: *n* = 11) (**A**) and CPH males (baseline: *n* = 16; week one: *n* = 11; week seven: *n* = 12) relative to SIH males (baseline: *n* = 12; week one: *n* = 11; week seven: *n* = 11) (**B**). Contrast maps show significant voxels at *p* < 0.005 with cluster size correction of *p* = 0.05. See Table 1 for brain regions and corresponding abbreviations.

### Sex differences in insula connectivity in the comorbid pain model

Since male rats exhibit less robust visceral hypersensitivity and referred pain-like behavior in the CPH model compared to females [19], we compared Ins FC between male and female CPH rats to dissect the brain mechanisms potentially driving sex differences in the development and progression of overlapping pain (Fig. 7; see Supplementary Fig. 1 for all significant brain regions). Compared to female CPH rats, male CPH rats demonstrate greater Ins FC with Pir1-3, En, orbital areas (L/V/VLO), PrL, motor areas (M1/2 and MAA), olfactory areas (AO and OB), BF, Acb, VP, CPu, insular areas (D/GI and AID/V), Cg1/2, Cl, IL, S1, RSD, Amy, Hyp, L/MPtA, CA3, and V2M. CPH females exhibit increased Ins FC to CPu, S1/2, CA1-3, and GP at baseline compared to their male counterparts.

**Figure 7.**
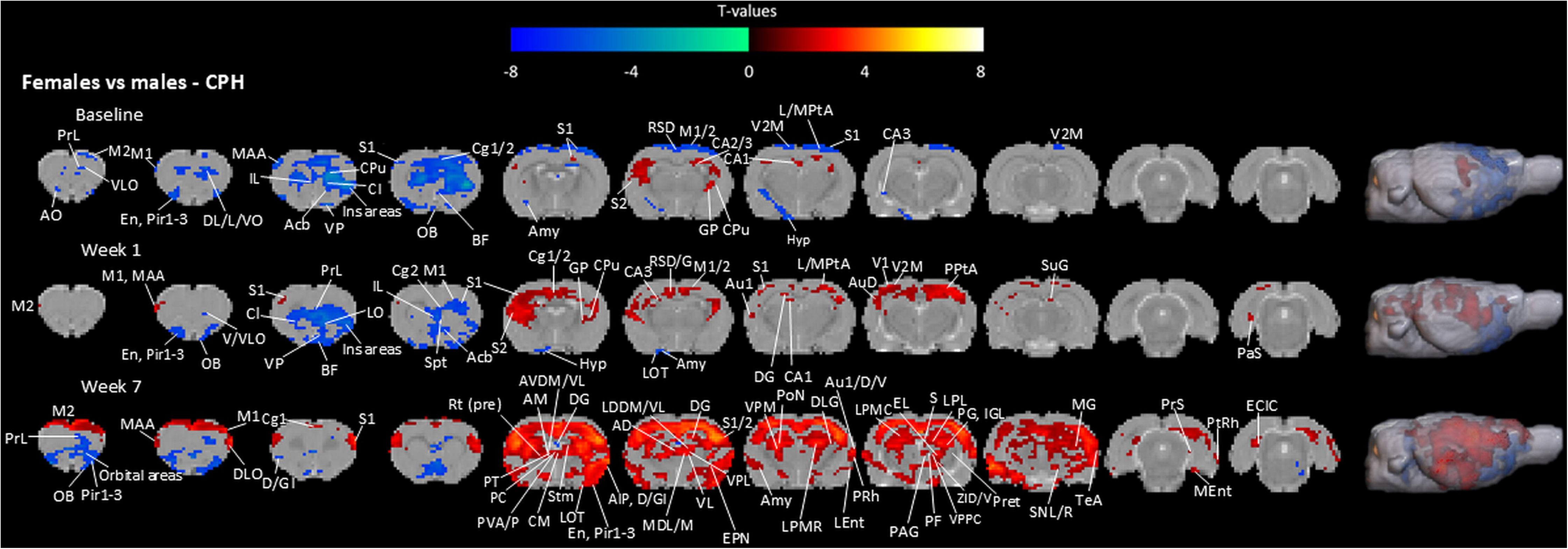
Effect of sex on insula-based connectivity in CPH model. Illustrated brain regions show increased and decreased connectivity in warm and cool colors, respectively, with Ins at baseline, week one, and week seven in CPH females (baseline: *n* = 16; week one: *n* = 18; week seven: *n* = 13) relative to CPH males (baseline: *n* = 16; week one: *n* = 11; week seven: *n* = 12). Contrast maps show significant voxels at *p* < 0.05 with cluster size correction of *p* = 0.05. See Table 1 for brain regions and corresponding abbreviations.

At week one, CPH males show increased Ins FC with Pir, orbital areas (L/V/VLO), En, olfactory areas (LOT and OB), BF, VP, CPu, PrL, insular areas (D/GI and AID/V), Cl, IL, Cg2, S1, Spt, Acb, M1, Hyp, and Amy. CPH females show increased Ins FC with motor areas (M1/2 and MAA), S1/2, Cg1/2, CPu, GP, DG, CA1/3, RSD/G, Au1/D, PPtA, V1/2, SuG, and PaS compared to their male counterparts at this time point.

At week seven, CPH males show greater Ins FC with olfactory areas (OB and AO), Pir1-3, orbital areas (DL/L/M/V/VLO), PrL, BF, VP, IL, D/GI, Acb, and DG. CPH females show vast differences in Ins FC, with increased Ins connectivity to the following brain regions: motor areas (M1/2 and MAA), DLO, S1/2, Cg1, CPu, Hyp, BF, Pir1-3, GP, Rt (pre), PT, AVDM, AVVL, PVA/P, PC, CM, AM, Cl, insular areas (D/GI and AIP), En, Amy, LOT, EPN, hippocampus (CA1-3 and DG), AD, LDDM/VL, VL, MDL/M, VPL, RSD/G, PRh, L/MEnt, Au1/D/V, DLG, VPM, PoN, LPMR, PtA, V1/2, PAG, Pret, S, PG, ZID/V, VPPC, PF, IGL, EL, LPL, LPMC/R, MG, SNL/R, TeA, PtRh, PrS, PaS, and ECIC. Week seven represents a time point at which visceral hypersensitivity and referred pain-like behavior has resolved in CPH males but not CPH females [19]. Accordingly, CPH females show a striking elevation in Ins-based connectivity to cortical areas and thalamic nuclei, including those involved in visceral pain and stress-induced anxiety, such as PV [48,50,53,85], suggesting that changes in Ins connectivity at various brain levels, notably the limbic system including the thalamus, may potentially contribute to sex differences in visceral hypersensitivity and referred pain-like behavior in the CPH model.

### Comorbid pain model time point differences reveal FC promoting persistent pain versus recovery

To better understand changes in Ins FC over time within our comorbid pain model, we compared Ins FC at baseline, week one, and week seven in CPH females and males (Fig. 8; see Supplementary Fig. 1 for all significant brain regions). In females (Fig. 8A), the following regions showed increased FC with Ins at week one compared to baseline: orbital areas (DL/L/V/VLO), PrL, motor areas (M1/2 and MAA), CPu, Cg1/2, Cl, GI, S1/2, Spt, hippocampus (CA1-3 and DG), RSD/G, L/M/PPtA, AuD, V1/2, S, and TeA. The following regions showed decreased connectivity to Ins in CPH females at week one compared to baseline: PAG, Pret, SuG, MG, PrS, ECIC, PtRh, PaS, and CIC. At week seven compared to baseline, CPH females showed increased connectivity to orbital areas (DL/L/V/VLO), PrL, motor areas (M1/2 and MAA), AO, Pir1/2, insular areas (D/GI and AID), Cg1/2, Cl, S1/2, CPu, BF, GP, Rt (pre), PT, AVVL, AM, hippocampus (CA1-3 and DG), Rn, VL, MDM, PVA/P, PC, CM, Amy, RSD/G, PRh, Au1/D/V, VPM, LDVL, PoN, ZID, LPMR, LPL, VPL, L/M/PPtA, V1/2, VPPC, EL, S, TeA, PtRh, and LEnt and decreased connectivity to SuG, PAG, PrS, ECIC, CIC, RSG, PtRh, and V1/2. This in accordance with our between-sex findings, as Ins increases connectivity with thalamic nuclei in CPH females at a time point where they exhibit continued visceral hypersensitivity and referred pain-like behavior compared to their male counterparts. Of note, this increased insular-thalamo connectivity does not appear at week one in CPH females compared to CPH males (Fig. 7), potentially reflecting a mechanistic switch subserving the transition from acute to chronic visceral hypersensitivity. Further supporting this trend, CPH females show increased Ins FC with the following thalamic nuclei at week seven compared to week one: Rt (pre), PT, AVDM/VL, PC, AM, LDVL, VL, MDM, PVA/P, CM, PoN, PF, and EL.

**Figure 8.**
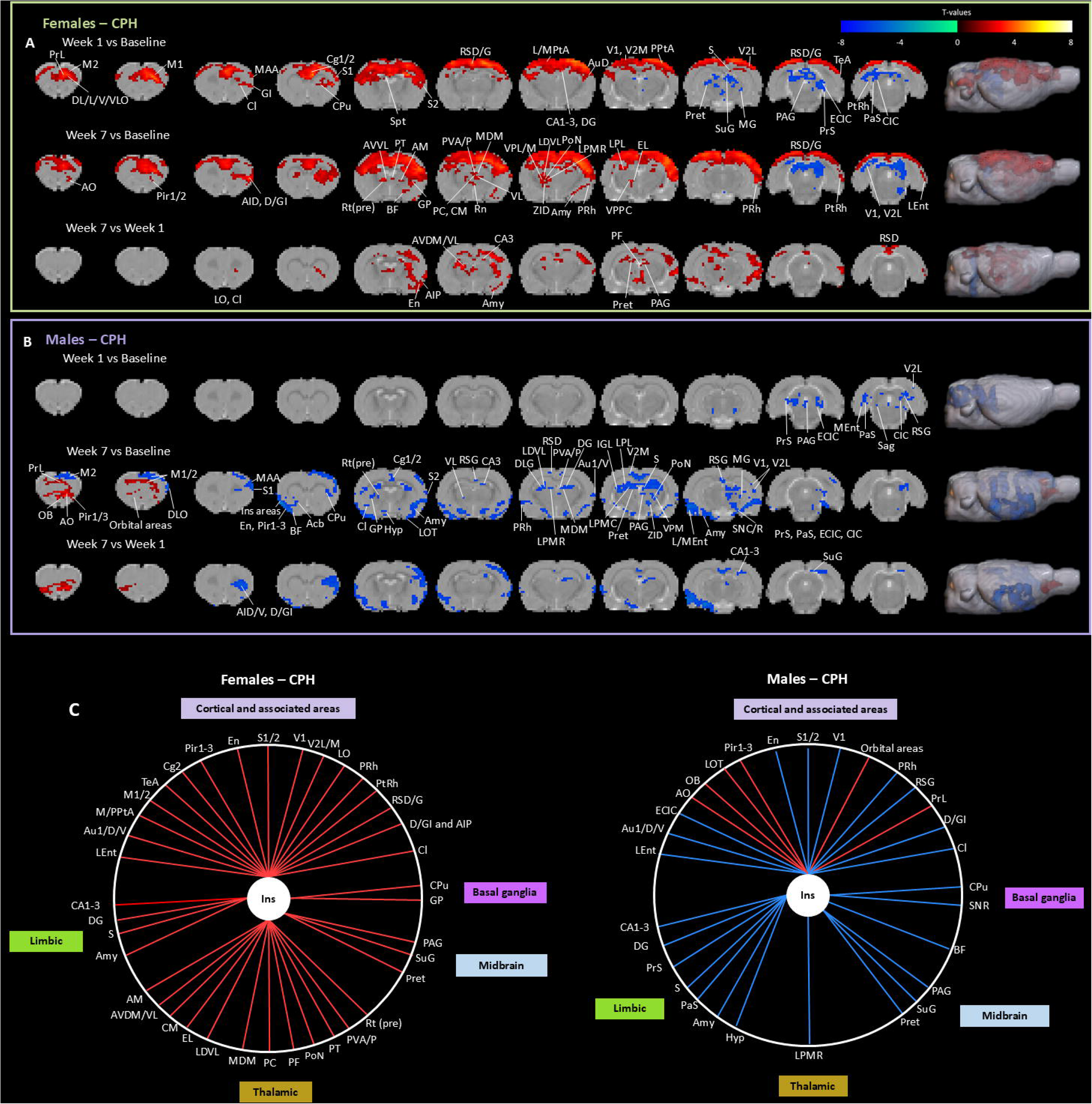
Changes in insula-based functional connectivity over time in CPH model. Illustrated brain regions show increased and decreased connectivity in warm and cool colors, respectively, with Ins at baseline, week one, and week seven in CPH females (baseline: *n* = 16; week one: *n* = 18; week seven: *n* = 13) (**A**) and CPH males (baseline: *n* = 16; week one: *n* = 11; week seven: *n* = 12) (**B**). Contrast maps show significant voxels at *p* < 0.05 with cluster size correction of *p* = 0.05. Week seven versus week one insula-based connectivity in CPH females and males is summarized in (**C**). Circle maps show brain regions with increased (red) or decreased (blue) connectivity. Brain regions are organized into associated networks. See Table 1 for brain regions and corresponding abbreviations.

At week one but not week seven, CPH males exhibit visceral hypersensitivity and referred pain-like behavior [19], so to understand the potential brain mechanisms contributing to the resolution of these pain outcomes in males, we compared Ins-based FC at baseline, week one, and week seven (Fig. 8B). Notably, a pattern of decreased connectivity with Ins emerged starting at week one compared to baseline, with weaker Ins FC to PAG, PrS, ECIC, CIC, PaS, MEnt, Sag, RSG, and V2L. In addition to the regions showing decreased connectivity to Ins at week one (with the exception of Sag), CPH males exhibited decreased connectivity at week seven compared to baseline with motor areas (M1/2 and MAA), DLO, S1/S2, BF, Pir1-3, Acb, insular areas (D/GI and AID), Cg1/2, En, Hyp, CPu, GP, Rt (pre), Cl, Amy, LOT, hippocampus (CA1-3 and DG), PRh, Au1/V, DLG, LDVL, MDM, PVA/P, V1, Pret, S, VPM, PoN, ZID, IGL, LPMC/R, LPL, LEnt, SNC/R, and MG. Increased connectivity with olfactory areas (OB and AO), Pir1/3, orbital areas (DL/L/M/V/VLO), PrL, and M1/2 appeared at week seven compared to baseline. Compared to week one, CPH males at week seven showed increased connectivity to olfactory areas (OB and AO), Pir1-3, orbital areas (DL/L/M/V/VLO), and PrL and decreased connectivity to BF, Pir1-3, insular areas (D/GI and AID), Cl, S1/2, CPu, En, Hyp, Amy, LOT, CA1/3, DG, PRh, Au1/D/V, LPMR, RSG, PAG, Pret, S, LEnt, SNR, SuG, PrS, V1, and ECIC. The clear decrease in Ins FC to thalamic nuclei at week seven compared to baseline and week one contrasts our findings in CPH females, highlighting that increased insular-thalamo connectivity may be a female-specific brain mechanism underlying prolonged visceral hypersensitivity and referred pain-like behavior in the CPH model. Overall, CPH males exhibit a trend of downregulating Ins connectivity (Fig. 8C), signaling that increased Ins connectivity may indeed contribute to worse pain outcomes in the CPH model.

## Discussion

The prevalence of COPCs, especially among women, has been increasingly recognized [54,69]. TMD and IBS are two commonly co-occurring chronic pain conditions that exhibit a female preponderance [75] and form positive feedback loops with psychological stress and mental health disorders, both of which share neural correlates with chronic pain [10,76]. Here, we aimed to characterize patterns of insular cortex (Ins) connectivity within our comorbid pain model, where stress following a priming injury (orofacial inflammation; TMD-like pain) generates *de novo* persistent visceral hypersensitivity and referred pain-like behavior (IBS-like pain) to a greater extent in female rats versus male rats [19]. Our previous study highlighted the Ins as a potential source of sexual divergence in visceral hypersensitivity, with CPH females showing increased Ins activity during colorectal distention at baseline and 1 week after stress compared to CPH males [19]. Using the Ins as a seed region, we observed disparate patterns of Ins connectivity by group, with a major finding being increased Ins connectivity to thalamic nuclei in CPH females at a time point where their visceral hypersensitivity remains—in direct contrast to the apparent resolution of visceral hypersensitivity in CPH males and SIH females.

The thalamus is a major supraspinal target of ascending pain pathways that conveys noxious information to cortical areas, which also encode the subjective aspects of pain [26]. As such, continuous noxious input can sensitize the neuraxis, including thalamo-cortical pathways, to incoming nociceptive signal (i.e., central sensitization). Likewise, conditions that feature aberrant central nervous system processing, such as stress [29], involve brain regions that overlap with the pain modulatory network, which can increase susceptibility to adverse pain outcomes including nociplastic pain [55]. The thalamus shares reciprocal connections with the Ins, a pain-processing brain hub consisting of posterior and anterior subdivisions, which are more involved in the sensorimotor and affective-motivational dimensions of pain, respectively [60,63]. CPH females demonstrated greater Ins functional connectivity (FC) with thalamic nuclei compared to CPH males at week seven (Fig. 7) and to their own baseline and week one connectivity (Fig. 8A and C). Among these thalamic nuclei, the paraventricular thalamic nucleus (PV) stands out, given its roles in visceral nociception and stress-induced anxiety-like behaviors [48,50,53,85]. In a model of colorectal visceral pain induced by neonatal colonic inflammation (NCI), optogenetically silencing PV terminals in the Ins attenuated visceral hypersensitivity, while optogenetically activating PV terminals elicited visceral hypersensitivity in naive mice [85]. The PV also responds to different stressors [8]. Specifically, the PV has demonstrated elevated neuronal activity days after predatory odor stress exposure [43] and increased alpha-2B adrenoceptor expression in the context of chronic psychosocial stress [36], highlighting stress-induced plasticity in the PV. Given that the PV seems to play a generalized role in stress, increased transmission to downstream targets, including the Ins, could potentially engender long-term network changes in chronic and subchronic stress paradigms. Of note, most of these studies did not include female rodents, so data is limited regarding sex differences in stress-induced changes in the PV and its downstream targets. However, employing an immobilization stress protocol, Ueyama et al. reported that estrogen-supplemented ovariectomized (OVX) female rats had greater c-Fos protein expression in the PV compared to OVX controls, suggesting a role for estrogen in the PV’s response to stress [81].

Another thalamic nucleus that responds to noxious visceral input and reciprocally connects with Ins is the mediodorsal thalamic nucleus (MD) [42,84], which showed greater FC with Ins at week seven in CPH females compared to CPH males (Fig. 7). Additionally, increased Ins FC to the ventromedial thalamic nucleus (VM) and amygdala (Amy) was observed at week seven in CPH females versus males (Fig. 7). The insula and amygdala share rich connections, and optogenetically silencing both Ins-basolateral amygdala and Ins-VM circuits induced analgesia and anti-depressant-like effects in a rodent model of neuropathic pain [15]. The role of the Ins and amygdala in aversion and aversion-associated states such as anxiety is conserved across species [28,35]; moreover, sexual dimorphism in amygdalar and thalamic mechanisms of pain chronification has been identified [77]. Besides PV, MD, and VM, we note increased Ins FC to various thalamic nuclei not directly linked to visceral pain, which could be attributed to dysregulation of affect/cognition-related insular-thalamic circuits and/or sensitization of intra-thalamic circuits, which may manifest physiologically as correlations in blood oxygen level dependent (BOLD) signal with Ins and/or behaviorally as the development and maintenance of referred pain, given that viscerosomatic convergence has been reported at the level of the thalamus [74].

Altogether, these findings suggest that sustained upregulation of Ins-thalamus connectivity may be a female-specific mechanism contributing to prolonged visceral hypersensitivity and referred pain-like behavior in the CPH model. Importantly, CPH males decrease Ins connectivity to thalamic nuclei, including PV and MD, at week seven compared to baseline (Fig. 8B), further supporting the idea that sensitization of the insular-thalamic network may potentially be a mechanistic switch driving the transition from acute to chronic visceral hypersensitivity and persistent referred pain-like behavior in females. Indeed, during colorectal distention, female rats have been shown to increase Ins-thalamus FC while males tend to downregulate this connectivity [15], suggesting sex-dependent differences in the processing of noxious visceral input. Sex-related differences in insular connectivity have also been reported in patients with IBS [37] and functional constipation [46].

Female CPH rats also displayed stronger Ins-cortical connectivity over time, particularly with sensorimotor areas, whereas males showed few changes in this network over time (Fig. 2). Female IBS patients with visceral hypersensitivity have shown increased resting FC of the posterior insula within the sensorimotor network, comprised of the motor cortices and supplementary motor area; this finding may reflect increased viscerosensory processing within the insula, promoting chronic visceral hypersensitivity [39]. Cortico-cortical modulation has been suggested to contribute to the attenuation of visceral pain in a model of colitis, where chemogenetic suppression of the midcingulate cortex reduced c-Fos expression in other cortical areas, including Ins [11]. Thus, cortico-cortical sensitization, including intra-insular circuits, could feasibly contribute to chronic visceral hypersensitivity in CPH females. Fig. 8C summarizes changes in Ins-based FC at week seven compared to week one in CPH females and males. There is a clear pattern of increased and decreased connectivity to Ins at various brain levels in CPH females and males, respectively, illustrating broad changes in Ins-based connectivity coincident with sexual divergence in visceral hypersensitivity and referred pain-like behavior.

Like CPH females, SIH females show extensive Ins-cortical and -thalamic connectivity over time; however, this trend dissipates in SIH females by week seven (Fig. 3), when they no longer exhibit visceral hypersensitivity, supporting the idea that sustained Ins hyperconnectivity to thalamic and cortical areas may contribute to chronic visceral hypersensitivity in the CPH model. When we compared SIH and CPH females directly, we provided more evidence to this point, with CPH females showing greater Ins connectivity with cortical areas such as S1/2, PRh, Cg1, visual areas (V1 and V2L/M), motor areas (M1/2 and MAA), and other Ins areas at week one and seven versus SIH females, who only showed decreased connectivity at week one with orbital areas (L/V/VLO) (Fig. 6A). Thus, stress may potentiate injury-induced Ins hyperconnectivity, namely with cortical areas, possibly leading to long-term central nervous system alterations that produce *de novo* nociplastic-like pain, whereas subchronic stress alone causes relatively transient neuronal and behavioral effects.

Given that our previous studies highlight sex differences in PAG activity [19–21], we unexpectedly observed similar patterns of Ins-PAG connectivity over time in CPH males and females. The PAG is a critical brain region in the descending pain modulatory system both in inhibitory and facilitatory capacities [79]. In fibromyalgia patients and healthy controls, PAG-to-insula connectivity was negatively correlated with the magnitude of conditioned pain modulation, a measure summarizing the efficiency of the descending pain modulatory system that has been associated with pain chronification [34]. Thus, decreased Ins-PAG connectivity observed in male and female CPH rats may reflect a reduction of pro-nociceptive signaling in the spinal cord; however, the PAG consists of different subcolumns not distinguished through our fMRI pipeline that play distinct roles in pain and stress-coping, so characterization of potential Ins-PAG involvement in the CPH model is needed. We also note unexpected differences in baseline Ins FC between groups, which may reflect individual variability and/or variability due to different sample sizes for each group.

We acknowledge that isoflurane anesthesia can alter brain activity. We administered a standard, low dose of isoflurane for a shorter duration than typical in-vivo approaches to minimize this effect while also avoiding potential stress that can be caused by awake protocols [57]. The use of various anesthesia protocols, including but not limited to isoflurane monoanesthesia, for rodent fMRI has been shown to alter functional connectivity. However, because of the limited information on the effects of combined anesthetics, such as low-dose isoflurane and medetomidine, on nociception and the continued use of low-dose isoflurane monoanesthesia in rigorous fMRI studies, we assert the validity of these findings having thoroughly considered and deliberated on alternative approaches. Finally, due to artifacts and image quality issues common in fMRI data collection, some animals were excluded from analysis at certain time points, as noted in the figure legends. Sample sizes did not decrease dramatically with subject exclusion, corresponding with subject number in other rodent longitudinal fMRI studies [18,38].

## Data availability

All data, including code, ROI, and the template brain are available upon request and in NeuroVault.

## Acknowledgments

This research was funded by the National Institutes of Health (National Institute of Dental and Craniofacial Research) grant number R01DE029074. The authors report no competing interests.

